# Quartet-based computations of internode certainty provide accurate and robust measures of phylogenetic incongruence

**DOI:** 10.1101/168526

**Authors:** Xiaofan Zhou, Sarah Lutteropp, Lucas Czech, Alexandros Stamatakis, Moritz von Looz, Antonis Rokas

## Abstract

Incongruence, or topological conflict, is prevalent in genome-scale data sets but relatively few measures have been developed to quantify it. Internode Certainty (IC) and related measures were recently introduced to explicitly quantify the level of incongruence of a given internode (or internal branch) among a set of phylogenetic trees and complement regular branch support statistics in assessing the confidence of the inferred phylogenetic relationships. Since most phylogenomic studies contain data partitions (e.g., genes) with missing taxa and IC scores stem from the frequencies of bipartitions (or splits) on a set of trees, the calculation of IC scores requires adjusting the frequencies of bipartitions from these partial gene trees. However, when the proportion of missing data is high, current approaches that adjust bipartition frequencies in partial gene trees tend to overestimate IC scores and alternative adjustment approaches differ substantially from each other in their scores. To overcome these issues, we developed three new measures for calculating internode certainty that are based on the frequencies of quartets, which naturally apply to both comprehensive and partial trees. Our comparison of these new quartet-based measures to previous bipartition-based measures on simulated data shows that: 1) on comprehensive trees, both types of measures yield highly similar IC scores; 2) on partial trees, quartet-based measures generate more accurate IC scores; and 3) quartet-based measures are more robust to the absence of phylogenetic signal and errors in the phylogenetic relationships to be assessed. Additionally, analysis of 15 empirical phylogenomic data sets using our quartet-based measures suggests that numerous relationships remain unresolved despite the availability of genome-scale data. Finally, we provide an efficient open-source implementation of these quartet-based measures in the program *QuartetScores*, which is freely available at https://github.com/algomaus/QuartetScores.

## Introduction

Recent advances in DNA sequencing technologies have greatly facilitated the generation of genome-scale data for phylogenetic inference in diverse groups of organisms, including fungi (e.g., Nagy et al. 2014; Shen et al. 2016), plants (e.g., Wickett et al. 2014; Yang et al. 2015), and animals (e.g., Jarvis et al. 2014; Misof et al. 2014). Incongruence (i.e., the presence of topological conflict) between individual gene trees in each one of these phylogenomic data matrices is the rule rather than the exception. The hundreds or thousands of genes examined in a study each yield their own distinct topologies (e.g., Song et al. 2012; Salichos and Rokas 2013; Zhong et al. 2013). The observed incongruence can be partly attributed to gene tree estimation errors caused by analytical reasons including insufficient information in the data, misspecification of evolutionary models, or inadequate tree search (Jeffroy et al. 2006; Kumar et al. 2012). On the other hand, the evolutionary histories of genes can also be genuinely different from each other and from the underlying species phylogeny due to biological processes such as incomplete lineage sorting, introgression, hybridization, and horizontal gene transfer (Maddison 1997; Slowinski and Page 1999; Degnan and Rosenberg 2009).

Given the prevalence of phylogenetic incongruence, its unequal distribution across branches of a phylogeny, and its key role in assessing the reliability of species tree inference (Salichos and Rokas 2013), it is important that our measures of incongruence are accurate. Salichos and colleagues recently developed several novel information theory-based measures to quantify incongruence among a set of “evaluation” trees (e.g., gene trees) with respect to the internodes (or internal branches) in a “reference” tree (e.g., the species tree) (Salichos and Rokas 2013; Salichos et al. 2014). In brief, for the bipartition defined by a given internode in the reference tree, its conflicting bipartitions are initially extracted from the evaluation tree set. Then, Shannon’s entropy (Shannon 1948) is calculated from the frequencies of occurrence (in the evaluation trees) of both the reference bipartition and the conflicting ones. In this way, the diversity and strengths of conflicting signals are integrated altogether as the degree of certainty (or uncertainty) about the phylogenetic relationship defined by the internode in the reference tree. The measures come in two flavors; the Internode Certainty (IC) score only takes into account the reference bipartition and the most prevalent conflicting bipartition, while the IC All (ICA) score also considers all other conflicting bipartitions that are sufficiently frequent.

The original IC/ICA scores are applicable only if all evaluation trees contain exactly the same taxa as the reference tree (Salichos et al. 2014). However, in phylogenomic studies, it is common that the sequences of many (or even most) genes are only available from taxon subsets. To meet the need to quantify incongruence in evaluation tree sets that contain partial trees, Kobert et al. (2016) developed mathematical approaches to adjust the frequencies of bipartitions from partial trees in the calculation of IC/ICA scores. Specifically, Kobert et al. (2016) developed three adjustment schemes that differ on how the frequency of a bipartition with missing taxa is corrected: 1) *Probabilistic* – the frequency of the incomplete bipartition is distributed equally to all possible comprehensive bipartitions (i.e., containing all taxa) that are compatible with it; 2) *Observed* – the frequency of the incomplete bipartition is distributed equally to only those compatible, comprehensive bipartitions observed in the reference and evaluation trees; and 3) *Lossless* – similar to *Observed*, but with the restriction that the comprehensive bipartitions also have to be mutually conflicting. Another approach similar to the *Lossless* adjustment scheme was also developed independently by (Smith et al. 2015).

IC and related measures are valuable and effective tools in revealing phylogenetic incongruence and have been quickly adopted in phylogenomic studies (e.g., Chen et al. 2015; Wang et al. 2015; Li et al. 2016; Shen et al. 2016; Chesters 2017; Krabberod et al. 2017; Leveille-Bourret et al. 2017), yet they still exhibit several practical and theoretical limitations. On the practical side, for data sets with high proportions of missing data (e.g., all genes trees are partial), the aforementioned adjustment schemes can considerably overestimate IC/ICA scores (Kobert et al. 2016). Additionally, alternative adjustment schemes might generate substantially different scores (Kobert et al. 2016) and it is often unclear which scheme is better. On the theoretical side, for the ICA measure, the exact number of conflicting bipartitions to be considered can only be determined *post hoc* from the evaluation trees (Salichos et al. 2014; Kobert et al. 2016), which might lead to unexpected behavior. To illustrate this point, consider the following example with one reference bipartition and two conflicting bipartitions. If we set their frequencies to 80%:10%:10%, 80%:15%:5%, and 80%:19%: 1%, the ICA scores would be 0.42, 0.44, and 0.51, respectively. That is, the ICA score of the internode increases as one of the conflicting bipartitions appears more frequently. However, if all 20% of the conflicting signal stems entirely from one bipartition (i.e., 80%:20%), then the ICA score drops again to 0.28. This is because the ICA score calculation now involves only two bipartitions instead of three, which changes the base of logarithm in Shannon’s entropy equation (Shannon 1948) from 3 to 2, thereby drastically lowering the score.

One potential solution to these practical and theoretical issues is to base the quantification of phylogenetic incongruence on quartets instead of bipartitions (see also (Pease et al. 2017)). Quartets (i.e., sets of four taxa) are the most basic unit of information in unrooted phylogenetic trees and have long been used in molecular phylogenetics for a wide range of purposes, including tree reconstruction (Strimmer and von Haeseler 1996; Chifman and Kubatko 2014; Avni et al. 2015; Mirarab and Warnow 2015), phylogenetic signal assessment (Strimmer and von Haeseler 1997; Nieselt-Struwe and von Haeseler 2001), and rogue taxon identification (Wilkinson 2006; Aberer and Stamatakis 2011). Several properties make quartets particularly attractive for quantifying IC. First, both the reference and evaluation trees can be decomposed into sets of induced quartets. Second, the quartet set of the reference tree is a superset of the quartet set of every evaluation tree. Therefore, both comprehensive and partial evaluation trees can be naturally compared with the reference tree at the quartet level without any further need for adjustment. In addition, evaluation trees with more missing taxa will contribute fewer quartets to the quantification (since the number of quartets contained in a tree is proportional to its number of taxa), providing a natural way to weigh evaluation trees of different sizes. Moreover, every quartet tree has a fixed number of three alternative topologies, hence two conflicting topologies will always be expected for every quartet topology in the reference tree regardless of the taxa present in the evaluation trees.

Here, we introduce three new quartet-based measures for quantifying incongruence among phylogenetic trees. Much like existing bipartition-based IC measures (Salichos et al. 2014; Kobert et al. 2016), the output of all three new measures are IC scores for all internodes in the reference tree, which reflect the degree of certainty of the bipartition defined by each internode. Using both simulated and biological data sets, we show that quartet-based and bipartition-based IC measures perform equally well in comprehensive trees and that quartet-based measures outperform bipartition-based ones on partial trees. Additionally, we establish the sensitivity of quartet-based IC measures to specific analytic challenges, such as the lack of phylogenetic signal and errors in reference trees. Finally, we applied our new measures on a comprehensive collection of empirical phylogenomic data sets and revealed prevalent phylogenetic incongruence in the eukaryotic tree of life. Overall, our results suggest that our newly developed quartet-based measures are useful for more accurately quantifying phylogenetic incongruence.

### Three New Quartet-Based Measures for Estimating Internode Certainty

All three quartet-based measures require as input a reference tree *T* and a set of evaluation trees *T̂*; only unrooted trees are considered. The taxon set of the reference tree S(*T*) should be equal to the union of the taxon sets of all evaluation trees S(*T̂*). All evaluation trees may have the same taxon set as S(*T̂*) (e.g., *T* and *T̂* are the bootstrap consensus tree and bootstrap replicate trees, respectively, from a single-gene phylogenetic analysis). Alternatively, the taxon sets of some or all evaluation trees
may be subsets of S(*T*) (e.g., *T* and *T̂* are the coalescent-based species tree and single-gene trees, respectively, from a phylogenomic analysis where some genes are missing from some taxa).

All three measures require the generation of a list of quartets induced by *T* and the occurrences of their alternative topologies in *T̂* (fig. 1A). Unresolved quartet topologies in polytomous evaluation trees are discarded. The three measures differ in whether all (or only a small fraction of all) possible quartets are used and how they are used, which in turn influences how the IC is calculated for each internode in *T* (fig. 1B-E).

**Figure 1.**
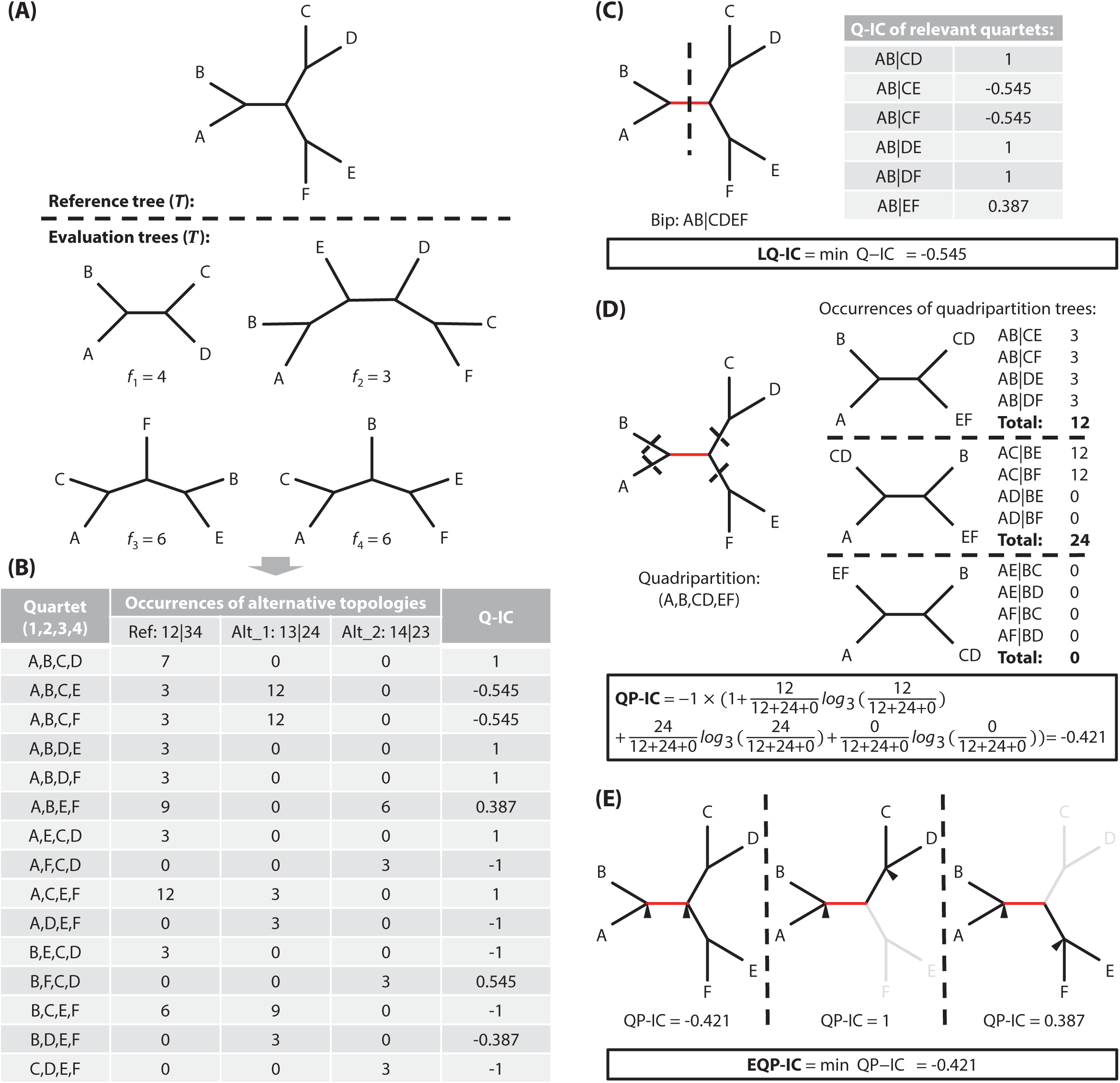
An example data set to illustrate the design and calculation of quartet-based IC scores. (A) The data set consists of a six-species reference tree *T* and an evaluation tree set *T̂*, each with a given frequency (shown along the respective tree topology). (B) The reference tree is decomposed into 15 quartets and the occurrences of their quartet tree topologies in the evaluation tree set are counted. This example focuses on the internode that separates (A, B) from (C, D, E, F). (C) The LQ-IC (Lowest Quartet IC) score of the internode is defined as the lowest IC score among all of its relevant quartets. (D) The QP-IC (Quadripartition IC) score of the internode is defined as the IC score of the quadripartition induced by it. (E) The EQP-IC (Extended Quadripartition IC) score of an internode is defined as the lowest QP-IC score among all of its relevant internal node pairs.

#### Measure 1: Lowest Quartet Internode Certainty (LQ-IC)

We define the LQ-IC of an internode as the lowest IC score among all of its relevant quartets (fig. 1B). Briefly, in a given unrooted tree, every internode defines a non-trivial bipartition, i.e., it divides the taxon set into two non-trivial subsets of taxa. We say a quartet *q* is relevant to an internode *i* if *q* consists of exactly two taxa from each of the two taxon subsets associated with *i*. For each internode *i* in *T*, we first identify the collection of all quartets (*Q*) that are relevant to *i*, and then calculate the IC score for each quartet *q* in *Q* based on the occurrences of its three possible topologies in *T̂* (*c*_1_, *c*_2_, and *c*_3_ for the alternative topologies *q*_1_, *q*_2_, and *q*_3_, respectively):

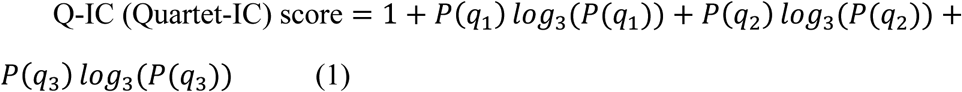

where *P*(*q*_1_) = *c*_1_/(*c*_1_ + *c*_2_ + *c*_3_), *P*(*q*_2_)) = *c*_2_/(*c*_1_ + *c*_2_ + *c*_3_), *P*(*q*_3_) = *c*_3_/(*c*_1_ + *c*_2_ + *c*_3_), and *P*(*q*_1_) + *P*(*q*_2_) + *P*(*q*_3_) = 1. This score equals 0 if *q* does not appear in any evaluation tree (i.e., *c*_1_ = *c*_2_ = *c*_3_ = 0). Also, we reverse the sign of the score if the topology of *q* induced by *T* is less frequent than any of the two alternative topologies.

Similar to IC/ICA scores, the Q-IC score can take values between −1 and 1: it approaches 1 when the reference quartet tree topology is much more prevalent than the other two alternatives, reflecting strong confidence in the reference internode; it becomes close to 0 when the three alternative topologies have similar frequencies, suggesting a high level of incongruence; and it gets near −1 when one of the conflicting topologies has a high frequency, indicating that the evaluation trees strongly contradict the internode present in the reference topology. A visualization of the Q-IC score against possible combinations of *P*(*q1*), *P*(*q2*), and *P*(*q3*) values is provided in supplementary fig. S1.

To obtain the LQ-IC score, we simply assign the lowest Q-IC score from *Q* to *i*:

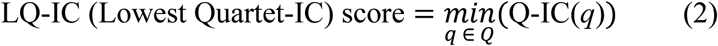

Since the calculation of the LQ-IC score for a given internode does not make any assumption about the topology on either side of *i*, LQ-IC can also be calculated for a reference tree *T* that contains polytomies (multifurcations); trees in the set of evaluation trees *T̂* may also contain polytomies.

#### Measure 2: Quadripartition Internode Certainty (QP-IC)

We define the QP-IC of an internode as the IC score of its induced quadripartition (fig. 1C). In a given unrooted binary tree, each internode connects two internal nodes (hereafter referred to as nodes) and divides the taxon set into four subsets (quadripartition). To determine the IC score of an internode, we assume that the four subsets have been correctly resolved and only consider the three possible topologies of the quadripartition. In other words, we consider the quadripartition as a “meta-quartet” whose leaves are the four subsets, and use the IC score of the “meta-quartet” tree as that of the internode.

For the quadripartition *p* induced by a given internode *i* in *T*, we calculate its IC score based on the occurrences of its three possible topologies in *T̂* (*c1*, *c2*, and *c3* for the alternative topologies *p1*, *p2*, and *p3*, respectively). We first identify the collection of all quartets (*Q*) that are relevant to *p*; we say a quartet *q* is relevant to a quadripartition *p* if *q* consists of exactly one taxon from each of the four taxon subsets associated with *p*. Each given quadripartition tree topology *t_p_*, induces a specific quartet tree topology *t_q_* for each *q* in *Q*, and the occurrence of *t_p_* is simply the sum of the occurrence of *t_q_* for all *q* in *Q*. We can then calculate the quadripartition IC score as:

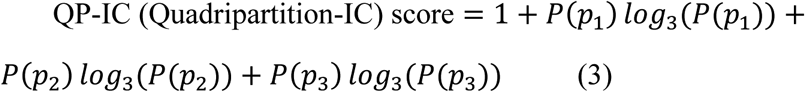

where *P*(*p*_1_) = *c*_1_/(*c*_1_ + *c*_2_ + *c*_3_), *P*(*p*_2_) = *c*_2_/(*c*_1_ + *c*_2_ + *c*_3_), *P*(*p*_3_) = *c*_3_/(*c*_1_ + *c*_2_ + *c*_3_), and *P*(*p*_1_) + *P*(*p*_2_) + *P*(*p*_3_) = 1. The equations (1) and (3) are almost the same, except that the former is calculated from a single quartet whereas the latter from a quadripartition. If the quadripartition tree topology induced by *T* is less frequent than any other alternative topologies, we reverse the sign of the QP-IC score. Unlike the LQ-IC score, the QP-IC can only be calculated for a binary reference tree *T.* The evaluation trees however may contain polytomies.

#### Measure 3: Extended Quadripartition internode certainty (EQP-IC)

In measure 2, we only consider quadripartitions induced by individual internodes, which means that only the quartets corresponding to neighboring nodes contribute to the IC score calculation. An alternative approach is to extend measure 2 to evaluate all possible pairs of nodes that include a given internode. Here, we define the EQP-IC score of an internode as the lowest IC score among all of its relevant node pairs (*N*) (fig. 1D); we say a pair of nodes *n* is relevant to an internode *i* if *i* is part of the path connecting *n*. Apparently, *N* includes both neighboring and non-neighboring node pairs. The IC score of a pair of neighboring nodes is simply its QP-IC score (see measure 2). In a given unrooted binary tree, every pair of non-neighboring nodes divides the taxon set into five subsets, four of which are directly associated with either node. We can therefore construct a “meta-quartet” from these four subsets and determine its QP-IC score also using measure 2. To obtain the EQP-IC score, we simply assign the lowest QP-IC score from *N* to *i*:

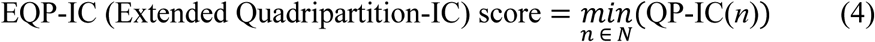

To calculate the EQP-IC score, the reference tree *T* must be binary, but the evaluation trees may be polytomous.

##### Example

To illustrate the calculation of our three quartet-based IC scores, we used an example data set consisting of a six-species reference tree *T* and an evaluation tree set *T̂* that includes one comprehensive and three partial trees, each with a given occurrence (shown along the respective tree topology; fig. 1A). In this example, we focus on the internode separating (A, B) from (C, D, E, F). In the first step, we decompose the reference tree into 15 quartets and, for each quartet, calculate the occurrences of the three possible topologies in the evaluation tree set (fig. 1B).

### LQ-IC score

The IC score of each quartet can be determined from the occurrences of its alternative topologies by equation (1) (fig. 1B). For instance, for the quartet (A, B, C, E), the reference topology (AB|CE) and the two alternative topologies (AC|BE) and (AE|BC) are respectively observed 3, 12, and 0 times in the evaluation trees.

Thus, for this quartet:

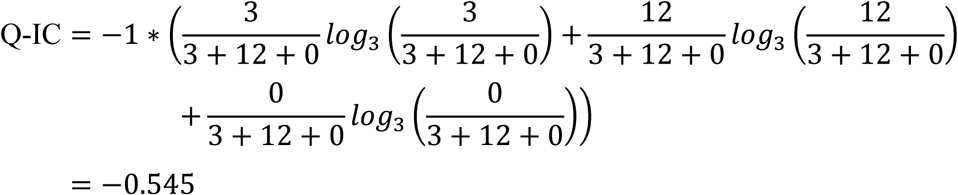

Note that the Q-IC score is negative since the reference quartet tree topology is less frequent than one of the conflicting topologies. Six of the 15 quartets are relevant to the internode of interest; therefore, the lowest Q-IC score among them (-0.545) is the LQ-IC score of the internode (fig. 1C).

### QP-IC score

The quadripartition induced by this internode is (A, B, CD, EF). The occurrence of each alternative quadripartition tree topology equals the sum of the occurrences of its induced quartet tree topologies (fig. 1D). For instance, the reference quadripartition tree topology, (A,B|CD,EF), induces four quartet trees: (AB|CE), (AB|CF), (AB|DE), and (AB|DF). Each quartet tree is observed three times in the evaluation trees, therefore the quadripartition tree has a total occurrence of 12. In the same way, the occurrences of the two conflicting topologies, (A,CD|B,EF) and (A,EF|B,CD), are found to be 24 and 0, respectively. Thus, for this quadripartition:

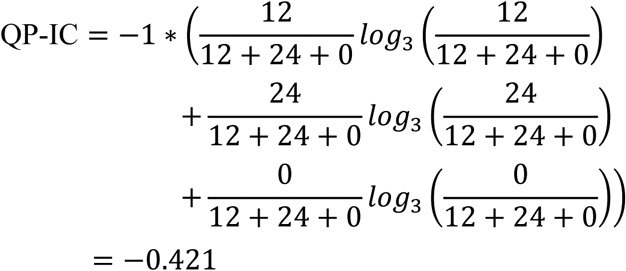

Note that the QP-IC score is negative since the reference quadripartition tree topology is less frequent than one of the conflicting topologies.

### EQP-IC score

The reference tree contains three node pairs that are relevant to the internode of interest (fig. 1E). The first one is the neighboring node pair that defines this internode itself, hence its QP-IC score equals that of the internode. The other two non-neighboring node pairs induce the quadripartitions (A, B, C, D) and (A, B, E, F), respectively, and their QP-IC scores are found to be 1 and 0.387 by following the same procedure described in the above section. Consequently, the lowest QP-IC score among the three node pairs (-0.421) is assigned to be the EQP-IC score of the internode.

## Results and Discussion

### Quartet-based and bipartition-based measures yield similar IC scores on comprehensive trees

We compared the performances of the quartet-based and bipartition-based measures on a simulated data set consisting of 50 101-taxon reference trees, each associated with 1,000 comprehensive evaluation trees (hereafter referred to as the “Original” data set; see Materials and Methods). The relative Robinson-Foulds (rRF) distance between the reference and evaluation trees range from 0.194 to 1 with a median value of 0.429. The three quartet-based IC scores are almost perfectly correlated with each other and the same is true for the bipartition-based IC/ICA scores (Spearman’s correlation coefficients ≥ 0.99 and p-values < 2.2×10^−16^ in all cases; supplementary fig. S2). Comparison of quartet-based IC scores with branch support values (measured by Gene Support Frequency (GSF); (Gadagkar et al. 2005)) and bipartition-based IC/ICA scores showed that quartet-based IC scores are strongly correlated with branch support values and bipartition-based IC/ICA scores (Spearman’s correlation coefficients ≥ 0.93 and p-values < 2.2×10^−16^ in all cases; fig. 2). At the same time, quartet-based IC scores tend to be more conservative than bipartition-based ones (fig. 2). Overall, the results suggest that the IC scores generated by quartet-based and bipartition-based measures are generally in agreement on data sets with only comprehensive trees.

**Figure 2.**
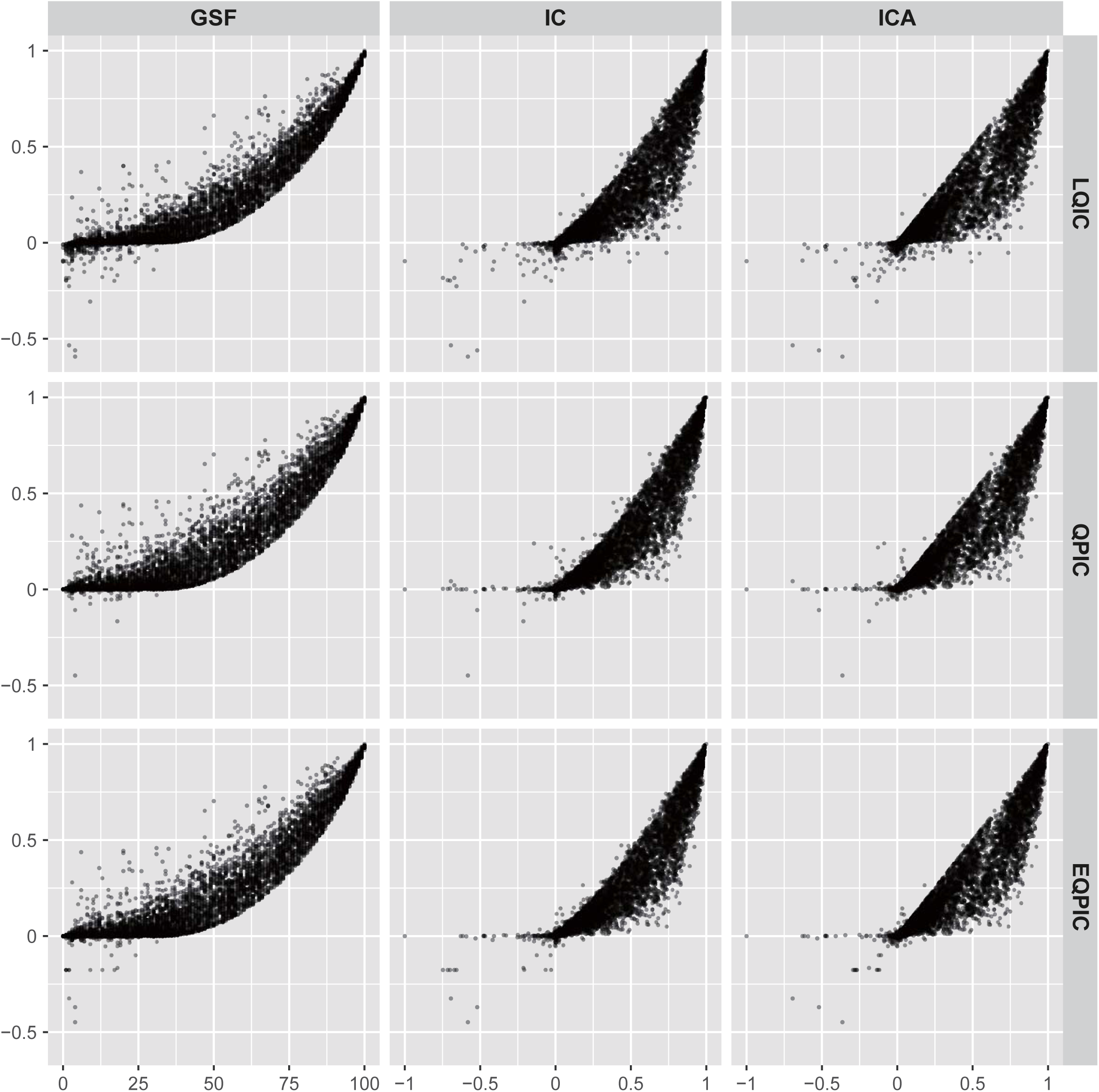
Strong positive correlations between IC scores generated by quartet-based measures and that generated by bipartition-based measures as well as the gene support frequency (GSF) values. The values shown were calculated on the Original data set containing only comprehensive evaluation trees. The spearman’s correlation coefficients range between 0.934 and 0.963.

### Quartet-based IC measures are more accurate on partial trees

Next, we compared the performance of quartet-based and bipartition-based IC measures on data sets with missing taxa. To that end, we constructed five additional data sets with partial trees – named L1, L2, L3, E1, and E2 – by pruning taxa from evaluation trees in the Original data set (see Materials and Methods for details). We note that: 1) the pruned taxa were randomly selected in L1, L2, and L3, while the patterns of missing taxa in E1 and E2 were sampled from empirical data sets; and 2) the degree of missing taxa increases in sequential order in L1, L2, and L3, while the degrees of missing taxa in E1 and E2 are comparable to those in L2 and L3, respectively (supplementary fig. S3). To examine the accuracy of each measure, we followed Kobert et al.’s (2016) suggestion that, on data sets with missing data, a more accurate measure should give scores that are closer to the ground truth. Here, we measured the accuracy by the Euclidean distance between the IC scores calculated on the Original data set (which we consider as the “truth”) and the pruned data sets. A smaller Euclidean distance indicates higher accuracy, and vice versa.

The results show that both quartet-based and bipartition-based measures exhibit similar accuracy when the proportion of missing data is low (data set L1; fig. 3A). However, while quartet-based measures remain highly accurate at medium to high proportions of missing data, the accuracy of bipartition-based measures decreases substantially (data sets L2, L3, E1, and E2; fig. 3A). One exception is the quartet-based LQ-IC measure, which also exhibits low accuracy on data set E2 (fig. 3A). Among the four bipartition-based measures, PIC and PICA – the two IC scores adjusted under the *Probabilistic* scheme – are consistently more accurate than LIC and LICA, which are adjusted under the *Lossless* scheme (fig. 3A). The same pattern is also apparent from the scatter plots of IC scores calculated on the Original and the pruned data sets (supplementary fig. S4).

**Figure 3.**
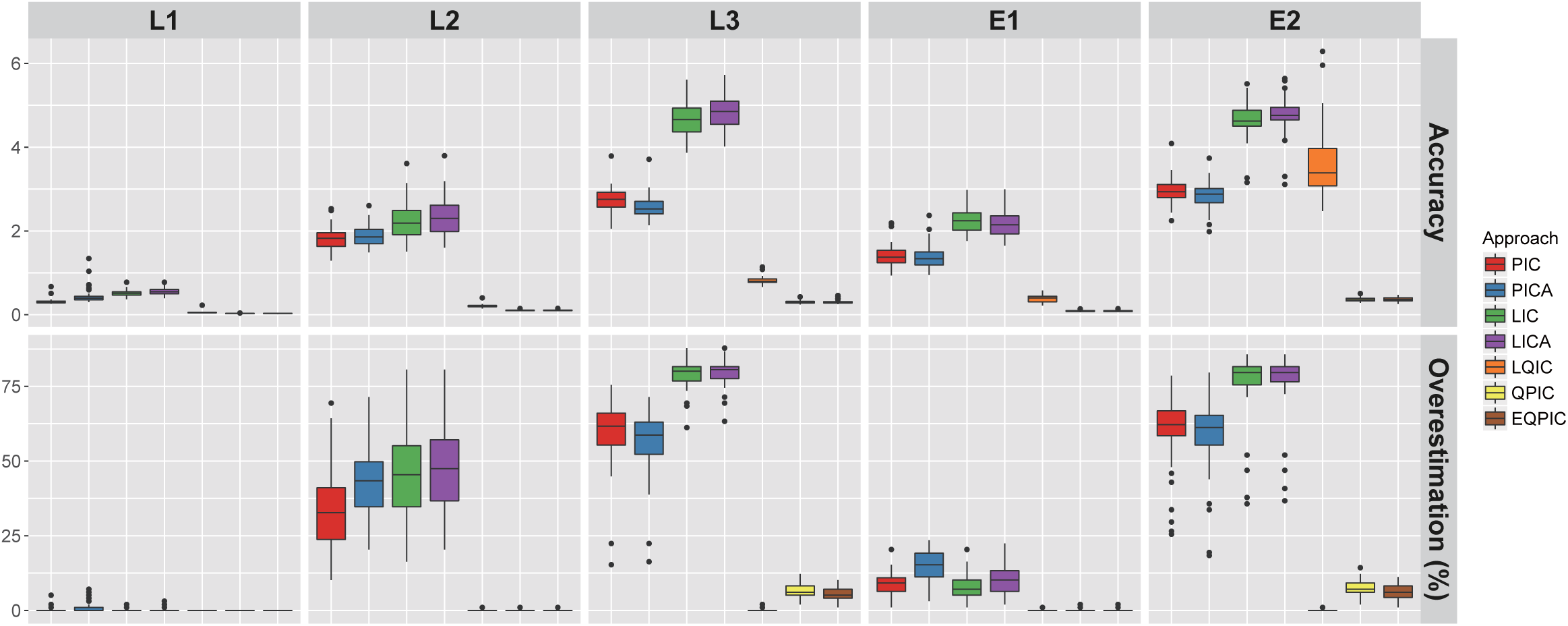
Quartet-based measures are more accurate on partial evaluation trees. (A) Euclidean distances between IC scores calculated from the Original data set which contain only comprehensive evaluation trees and those calculated from data sets L1-3 and E1-2 which contain partial evaluation trees. (B) Fractions of internodes for which the IC scores were overestimated (0.05 unit higher) on data sets L1-3 and E1-2 compared with the Original data set. The boxplots depict, for each measure, distribution of Euclidean distance (A) or overestimated fraction (B) values from 50 replicates.

For each measure, we also calculated the fractions of internodes for which the IC scores were overestimated on the pruned data sets. We observed a trend very similar to that found in the accuracy assessment. While quartet-based measures exhibit very low levels of overestimation, bipartition-based measures tend to overestimate IC scores on pruned data sets (fig. 3B). In particular, bipartition-based IC scores are overestimated for most of the internodes at high proportions of missing data (data set L3 and E2; fig. 3B). Altogether, these results suggest that quartet-based IC measures are more accurate than bipartition-based measures on partial evaluation trees.

We further compared the quartet-based and bipartition-based IC measures on two empirical data sets previously analyzed in Kobert et al. (2016), namely a 23-taxon yeast data set containing 1,275 comprehensive gene trees and 1,219 partial gene trees, and an avian data set containing 500 comprehensive gene trees and 1,500 partial gene trees. IC scores calculated from all gene trees were compared with the scores calculated from either only the comprehensive trees or only the partial trees. Here again, we found that the Euclidean distances for quartet-based measures are considerably lower than the distances for bipartition-based measures in all comparisons (fig. 4; see supplementary fig. S5 for detailed scatter plots of IC scores), suggesting that quartet-based IC scores are more consistent with the IC obtained from the comprehensive trees.

**Figure 4.**
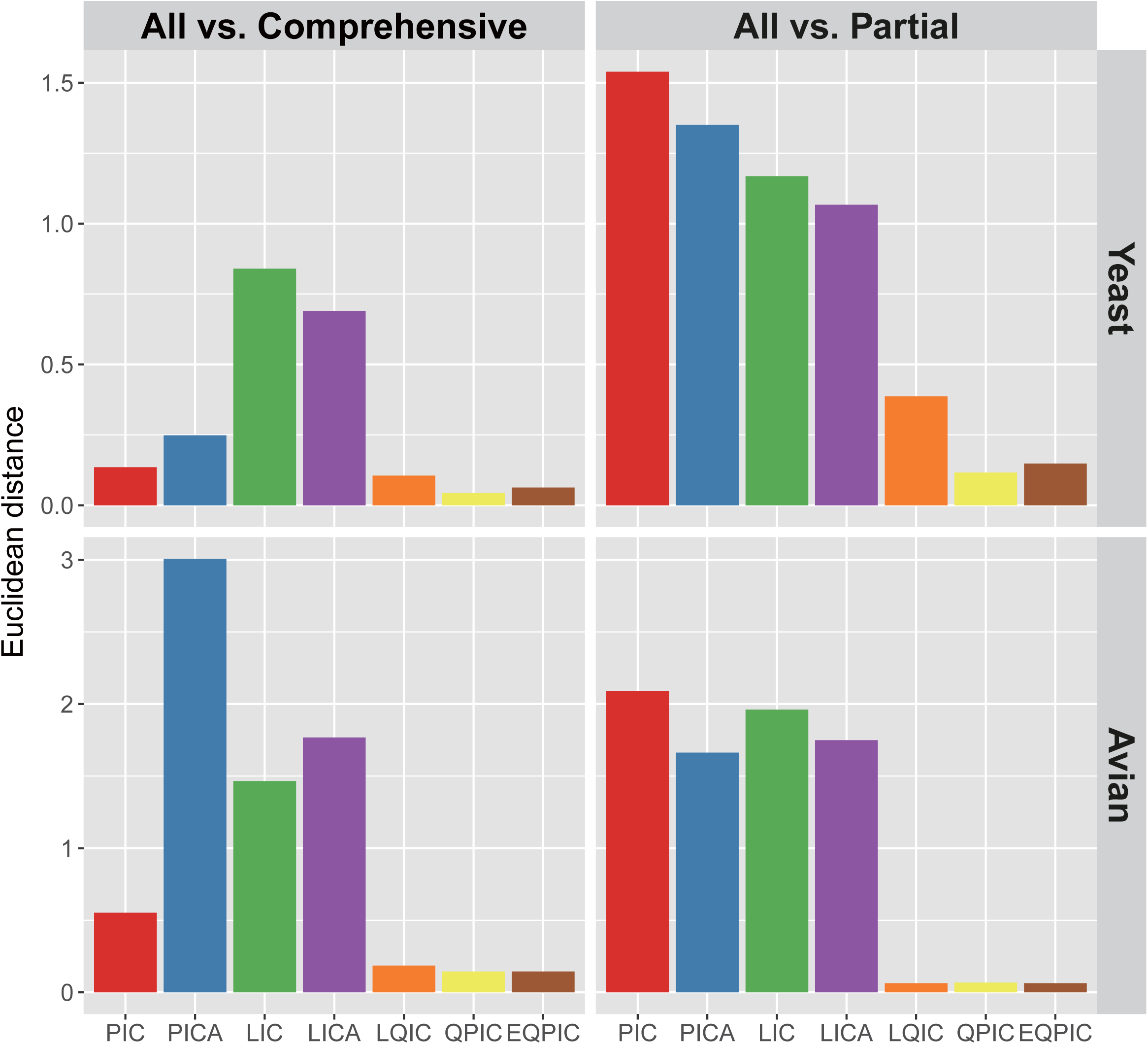
Euclidean distances between IC scores calculated on only comprehensive evaluation trees or only partial evaluation trees versus the scores calculated on all trees. The results of the (A) yeast and (B) avian empirical data sets are shown.

### The robustness and vulnerability of IC measures to specific analytical challenges

To investigate the potential strengths and/or limitations of different IC measures, we next assessed their performance under three analytical challenges, namely lack of phylogenetic signal, high proportions of missing data, and errors in reference trees.

#### Lack of phylogenetic signal

In the aforementioned analysis of the empirical avian data set, we observed that, for some internodes, the quartet-based IC scores are around 0, a value indicative of two nearly equally supported conflicting resolutions, whereas the bipartition-based IC scores are near or at −1, a value indicative of the presence of a strongly supported conflicting bipartition (supplementary fig. S5). Closer examination of the underlying bipartition frequencies at these internodes revealed that none of the conflicting bipartitions is supported (supplementary table S1). For instance, for multiple internodes, the reference bipartition and the most prevalent conflicting bipartition have frequencies of 0 and 0.034, respectively. This suggests that bipartition-based IC measures might report strong support for a conflicting bipartition when in reality there is little phylogenetic signal.

To test this behavior of bipartition-based IC scores further, we devised a “random evaluation tree” test where we used completely random evaluation tree topologies in the Original, E1, and E2 data sets (see Materials and Methods). Thus, in principle, the evaluation tree sets should provide no support to any particular relationship and the IC scores for all internodes should be near or at 0. The results of this test show that bipartition-based IC scores (except for PICA scores) are indeed heavily skewed toward −1 (fig. 5). Interestingly, LIC and LICA scores show much less bias on data set E2 (high proportion of missing data) than on E1 (medium proportion of missing data) (fig. 5B and C.). In contrast, quartet-based measures in general show a more robust performance; the only exception is LQ-IC, which generates artificially low scores at high proportion of missing data (data set E2; fig. 5C).

**Figure 5.**
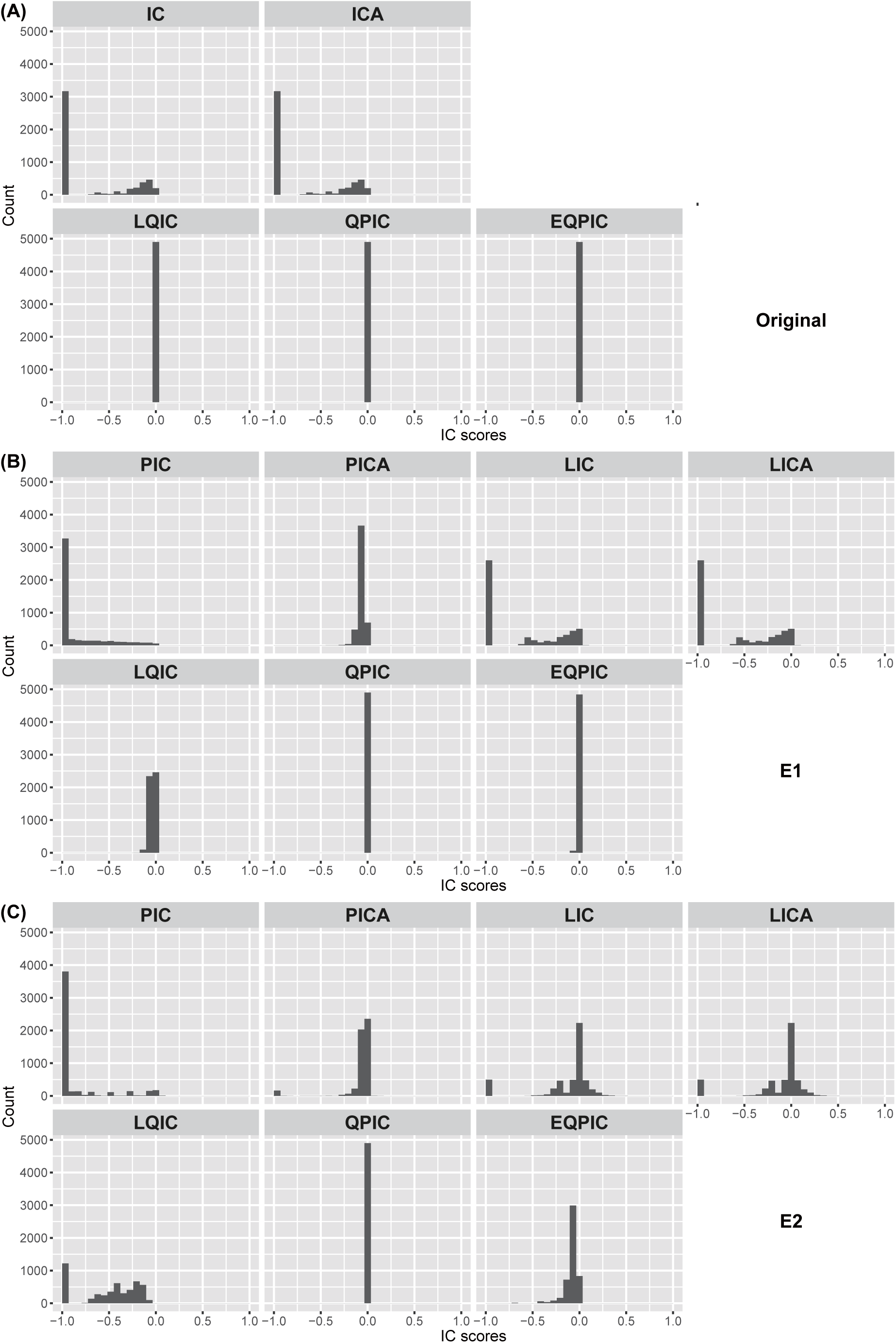
Bipartition-based IC scores tend to generate artificially low scores in the lack of phylogenetic signal. The results of the “random evaluation tree” test on data sets (A) Original, (B) E1, and (C) E2 are shown. The histograms indicate, for each measure, the distribution of IC scores calculated using randomized evaluation trees.

#### High proportion of missing data

In the accuracy assessment on simulated data, we noted that many internodes have positive LQ-IC scores in the Original data set, but exhibit scores of −1 on E2 (supplementary fig. S3). One possible explanation for the impaired performance of LQ-IC on data set E2 is that, due to the high proportion of missing data, some quartets are present in but a few evaluation trees. Consequently, the IC scores of these quartets resemble random noise rather than reflecting the level of incongruence, and the LQ-IC scores of some internodes might be driven by the quartets that get low IC scores by chance. We investigated the IC scores of individual quartets in the six simulated data sets and found that, indeed, the quartets that have QIC scores near −1 in E2 all occur infrequently (supplementary table S2). In addition, the Q-IC scores of E2 deviate from the Original data set much more than in other data sets (supplementary fig. S6).

#### Errors in reference tree

One important assumption underlying the design of the QP-IC and EQP-IC scores is that the four subsets of taxa around a given internode are correctly resolved (referred to as the “locality assumption” in (Sayyari and Mirarab 2016)). To test the performance of QP-IC and EQP-IC when the locality assumption is violated and also the performance of other IC measures on reference trees containing incorrect relationships, we devised an “altered reference tree” test. In this test, varying degrees of errors were introduced into the reference trees in the Original, E1, and E2 data sets; the corresponding rRF distances between original and altered reference trees range between 0.1 and 1 (see Materials and Methods).

We examined the IC scores of the bipartitions that are only present in the altered reference trees but not in the original ones. Since the vast majority of internodes in the original reference trees have positive IC scores, the bipartitions introduced by the alterations are expected to be contested by other, higher-frequency bipartitions (supplementary fig. S3). Indeed, most of the introduced internodes have negative LQ-IC and EQP-IC scores as well as negative bipartition-based IC scores in the absence of missing data (data set Original; fig. 6A). Conversely, the QP-IC scores are positive for a considerable fraction of these introduced internodes (fig. 6A). Interestingly, all the IC measures generate lower scores on altered reference trees that are more dissimilar to the original trees than on altered reference trees that are more similar to the original trees (fig. 6A). The same patterns are also observed on data set E1, which has a medium proportion of missing data (fig. 6B). However, at high proportion of missing data (data set E2), bipartition-based measures produce positive scores for even more incongruent internodes than QP-IC (fig. 6C). In contrast, the performances of LQ-IC and EQP-IC are consistent at all different proportions of missing data (fig. 6).

**Figure 6.**
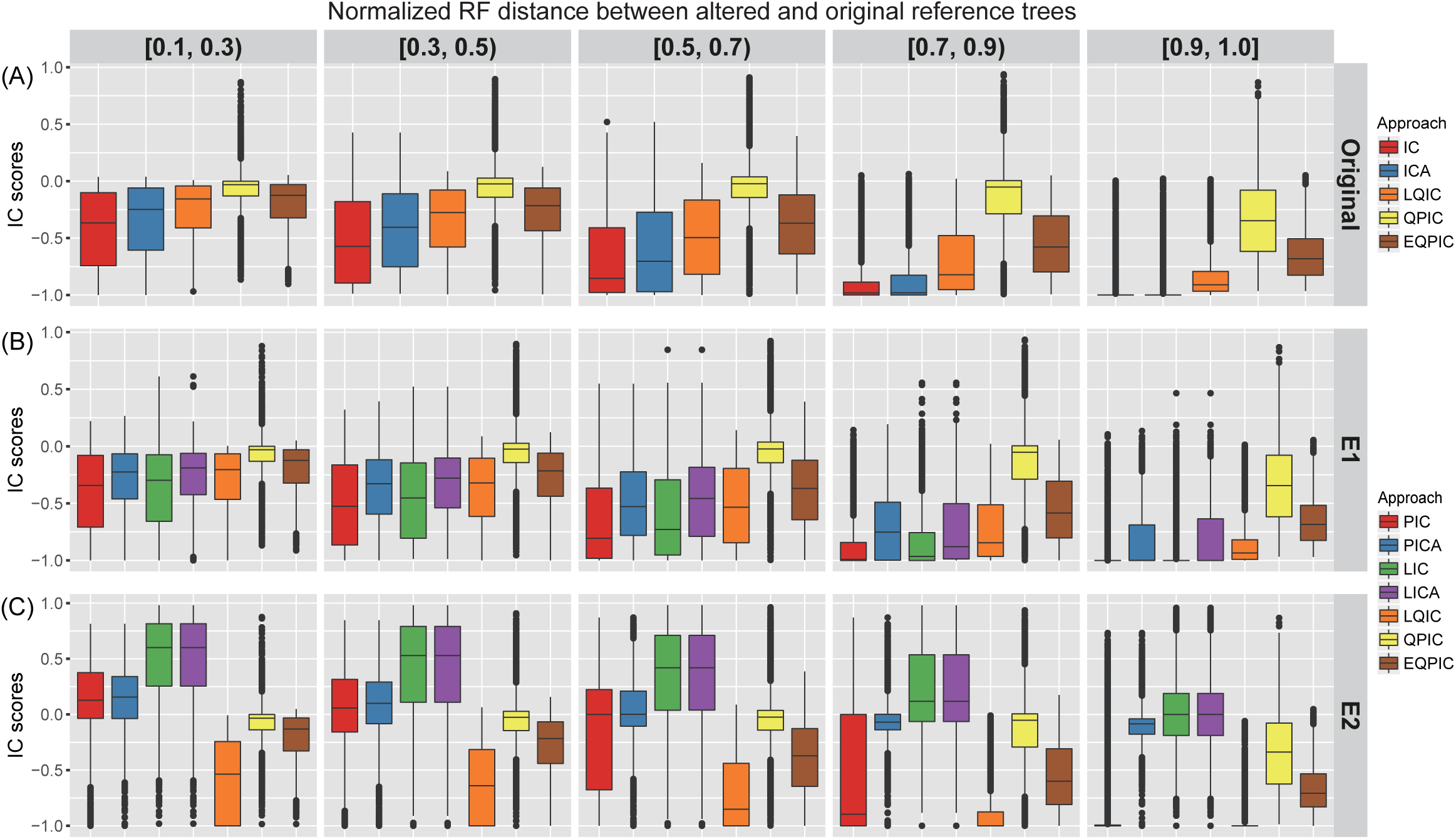
The robustness of quartet-based and bipartition-based IC measures to errors in reference trees. The results of the “altered reference tree” test on data sets (A) Original, (B) E1, and (C) E2 are shown. The boxplots indicate, for each measure, the distribution of IC scores for bipartitions that are only present in the altered reference trees.

These results suggest that the violation of the locality assumption can often lead to inflated QP-IC scores. The related EQP-IC measure seems robust to such violations; the locality assumption might be relaxed due to the consideration of other non-neighboring node pairs in the EQP-IC measure. Bipartition-based IC measures perform generally well except at high proportion of missing data. The reason might be that, as has been shown earlier in this study, bipartition-based measures tend to overestimate IC scores when the proportion of missing data is high (fig. 3B).

##### Analysis of empirical phylogenomic data sets

Having validated their performance, we applied our quartet-based IC measures along with previous bipartition-based measures to a comprehensive collection of 15 empirical data sets from recent phylogenomic studies of fungi (Nagy et al. 2014; Shen et al. 2016), plants (Wickett et al. 2014; Xi et al. 2014; Yang et al. 2015), and animals (Song et al. 2012; Jarvis et al. 2014; Misof et al. 2014; Borowiec et al. 2015; Chen et al. 2015; Prum et al. 2015; Struck et al. 2015; Whelan et al. 2015; Tarver et al. 2016) (supplementary table S3). We calculated IC scores for each data set using the coalescent-based species tree as the reference tree and the single-gene trees (from which the species tree was estimated) as evaluation trees (see Materials and Methods).

We first describe the results from quartet-based IC measures. The data sets display varying degrees of phylogenetic incongruence (fig. 7); the average LQ-IC score of each data set ranges between −0.709 and 0.494, while the average QP-IC and EQP-IC scores range between 0.131 and 0.532. For most data sets, the average IC scores are higher than the median values, suggesting that many internodes may have low IC scores. In fact, more than one third of all internodes in the 15 data sets have LQ-IC, QP-IC, and EQP-IC scores between −0.1 and 0.1 (fig. 7; supplementary table S4); quartet-based IC scores in this range correspond to very weak support for the corresponding internodes in the reference topologies and are suggestive of highly uncertain relationships (supplementary fig. S1). At the level of individual data sets, 9 of the 15 data sets show the same pattern (fig. 7; supplementary table S4).

**Figure 7.**
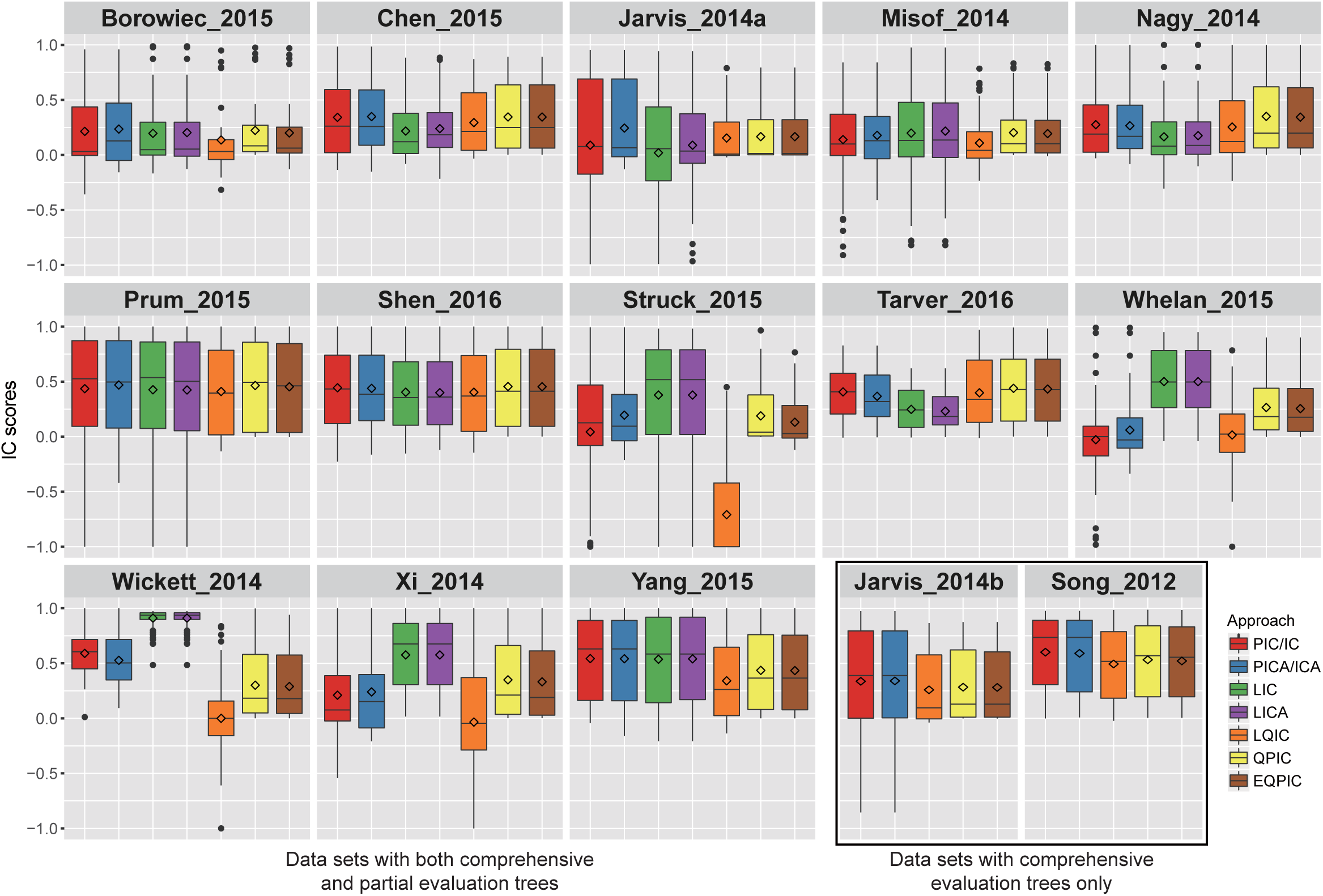
Analysis of 15 empirical phylogenomic data sets using quartet-based and bipartition-based IC measures. The boxplots indicate, for each measure, the distribution of IC scores for all internodes in each data set. The horizontal line and diamond in each boxplot indicate the median and mean of the IC scores, respectively.

Notably, the quartet-based IC scores almost never show values near −1; the minimum QP-IC and EQP-IC scores among all internodes in the 15 data sets are - 0.004 and −0.128, respectively (fig. 7; supplementary table S4), suggesting that there are no strongly supported conflicting relationships. The more conservative LQ-IC measure also generates minimum scores in the range between −0.316 to −0.012 except for four data sets (Struck_2015, Whelan_2015, Wickett_2014, and Xi_2015) in which some internodes have scores of −1 (fig. 7; supplementary table S4). However, the LQ-IC scores are likely biased in these four data sets due to high proportions of missing data (supplementary table S4). Overall, our results on these empirical data sets suggest that certain relationships may remain unresolved even though genome-scale data amounts have been analyzed (see also (Shen et al. 2017)).

Bipartition-based measures give rise to similar median IC scores as quartet-based ones for most data sets (fig. 7). However, the scores at the level of individual internodes often differ considerably between the two types of measures (supplementary fig. 7). Additionally, bipartition-based measures also produce scores near or at −1 for several data sets, which is – as discussed above – likely an artifact observed due to the lack of phylogenetic signal (supplementary table S5).

Furthermore, the scores generated by alternative adjustment schemes (PIC/PICA versus LIC/LICA) show substantial discrepancies on the four data sets where all evaluation trees are partial (Struck_2015, Whelan_2015, Wickett_2014, and Xi_2015; fig. 7), consistent with the observation of Kobert et al. (2016) that the inclusion of comprehensive evaluation trees is critical to the performance of bipartition-based measures.

##### Related measures of branch support or phylogenetic incongruence

Besides the IC scores discussed above, there also exist other related measures of branch support or phylogenetic incongruence. Importantly, our QP-IC measure is based on the idea of comparing alternative arrangements of the four clades around a given internode. The same concept has been previously applied in three fast likelihood-based branch support tests, including the approximate likelihood-ratio (aLRT) test (Anisimova and Gascuel 2006), the Shimodaira–Hasegawa variant of aLRT (Guindon et al. 2010), and the Bayesian variant of aLRT (Anisimova et al. 2011). All three tests evaluate the likelihood scores of the three possible topologies for each internode by applying the two possible NNI (nearest neighbor interchange) moves without changing the subtrees defined by the internode. The measures can therefore be computed efficiently.

The local posterior probability (LPP) is another fast method for local branch support (Sayyari and Mirarab 2016). Similar to QP-IC, it first calculates the quartet-based frequencies of alternative quadripartition topologies around a given internode in the species tree from a set of gene trees. The probability that the quadripartition is present in the true species tree (i.e., the LPP), is then estimated under the multispecies coalescent (MSC) model. By invoking the MSC model, LPP explicitly accounts for incomplete lineage sorting which is an important source of species tree-gene tree discordance. In comparison, neither our quartet-based nor the original bipartition-based IC measures make any assumption on the underlying causes of incongruence. Therefore, these measures are broadly applicable to measuring the level of incongruence in any data type (e.g., a maximum-likelihood gene tree as the reference tree and the corresponding bootstrap replicate trees as evaluation trees).

In parallel to this work, Pease et al. (2017) developed the Quartet Sampling (QS) measure, which integrates several features of the above mentioned methods; it is a likelihood-based (like aLRT tests), quartet-based (like LPP), and entropy-based measure of phylogenetic incongruence (like IC measures). The major distinction between QS and our QP-IC measure is that, in QS, quartets are randomly sampled and the three alternative topologies for each quartet are evaluated independently under the ML criterion, whereas in QP-IC, all quartet tree topologies are extracted from already estimated evaluation trees. Accordingly, QS requires only the reference tree but can only be applied to a single data matrix; on the other hand, our quartet-based measures require pre-estimated evaluation trees, but can be used for both single data matrix analysis (on bootstrap replicate trees) and for coalescent analysis (on single-gene trees).

Further studies are needed to compare the performances of QS and our quartet-based IC measures on phylogenomic data sets. On one hand, the evaluation of quartet tree topologies in the QS measure might be sensitive to phylogenetic artefacts such as long-branch attraction (Ranwez and Gascuel 2001). On the other hand, the performance of quartet-based IC measures can be impaired by inaccurate gene tree estimation when the numbers of taxa become high and the lengths of single-gene alignments become short. Nevertheless, the two types of measures can complement each other and their joint usage in phylogenomic studies will likely yield a more comprehensive understanding of phylogenetic incongruence in genome-scale data sets.

##### Implementation

We have implemented the three quartet-based IC measures in the program *QuartetScores*, which is freely available as open source code at https://github.com/algomaus/QuartetScores (last accessed July 24, 2017). The calculation of quartet-based IC scores requires counting the occurrences of all quartet tree topologies in the evaluation tree set, a challenging task since the number of possible quartets grows exponentially with the number of taxa. Therefore, we devised two algorithms for quartet counting: one is more time-efficient by storing each quartet topology separately in a lookup table; the other is more memory-efficient by grouping different topologies of a quartet together and using a more complicated indexing function. The program will automatically decide which algorithm to use based on the data set size. A full description of the algorithms for counting quartets and computing quartet-based IC scores is provided in the supplementary text.

## Materials and Methods

### Simulated data sets

We used a simulated data set from Mirarab and Warnow (2015) (referred to as the “Original” data set in our study) to evaluate the performance of our quartet-based IC scores. This Original data set contains 50 sets of trees, each of which has one species tree and 1,000 estimated gene trees. In brief, for each set, a 101-taxon species tree was first simulated according to the Yule process with a speciation rate of 10^-6^ per generation and a tree length of 2 million generations. Then, 1,000 gene trees were simulated on the species tree under the multiple-species coalescent model. For each simulated gene tree, a gap-free nucleotide alignment was simulated under the GTR+GAMMA model, and FastTree 2 was used to infer a maximum-likelihood gene tree. The simulated species trees and their corresponding estimated gene trees were downloaded from https://www.cs.utexas.edu/~phylo/datasets/astral2 and used in this study (last accessed July 24, 2017).

All gene trees in the Original data set are comprehensive. To further examine the performance of quartet-based IC scores on partial trees, we generated five additional data sets by pruning taxa from gene trees in the Original data set. The taxon-pruning was conducted in two different ways. For three of the five data sets (P1, P2, and P3), taxa were pruned randomly and, for each gene tree, the number of taxa to prune was drawn from a log-normal distribution (truncated on the right at 97 to ensure that pruned trees have at least four taxa). The three data sets were generated by using lognormal distributions with mean values of ln 1, ln 10, and ln 100, respectively, corresponding to low, medium, and high proportions of missing data.

For the other two data sets (E1 and E2), the patterns of missing taxa were sampled from empirical data sets to better approximate real data conditions. For instance, E1 was generated by using the 144-taxon, 1478-gene empirical data set from Misof et al. (2014) as template as follows: first, 101 taxa and 1,000 gene trees were randomly selected from the empirical data set and randomly paired with the taxa and gene trees in the Original data set; then, for each taxon *t* and gene tree *g* selected from the empirical data set, if *t* is missing from *g*, the paired taxon *t*’ was pruned from the paired gene tree *g*’ in the simulated Original data set. This procedure was performed independently for each of the 50 sets of trees in the Original data set. The data set E2 was constructed in the same way based on the 103-taxon, 844-gene empirical data set from Wickett et al. (2014) (note that some genes were sampled twice from the empirical data set, since it has less than 1,000 genes). All reference and evaluation trees used in this study, as well as the results of IC analyses are available from the figshare repository (https://figshare.com/s/2e44121d0c3ff60004e3; last accessed July 24, 2017).

### Empirical data sets

Two collections of empirical data sets were analyzed in this study. We first compared the performance of quartet-based and bipartitions-based IC scores on the 23-taxon yeast data set and the 48-taxon avian data set, which were used in Kobert et al. (2016) to evaluate various adjustment schemes of the original IC/ICA scores. The two data sets were originally published in Salichos and Rokas (2013) and Jarvis et al. (2014), respectively. Gene trees and species trees in these two data sets were downloaded from https://github.com/Kobert/ICTC (last accessed July 24, 2017). We also calculated various IC scores for 15 data sets from 14 recently published phylogenomic studies (see the details of the data sets in supplementary table S3) to investigate the pattern of phylogenetic incongruence across the eukaryotic tree of life. These data sets have been re-analyzed systematically in a recent study using several fast maximum likelihood-based phylogenetic inference programs (Zhou et al. 2017). The resulting best-scoring gene trees and coalescent-based species trees (estimated from the gene trees using ASTRAL) were downloaded from https://goo.gl/VkMTa4 (last accessed July 24, 2017).

### Calculation of branch support and IC scores

For a given reference tree and evaluation tree set, the gene support frequencies for internodes in the reference tree were calculated using RAxML v8.2.10 with the “-f b” option. Similarly, the bipartition-based IC scores were calculated by using RAxML with the “-f i” option. The original IC/ICA scores were reported if all evaluation trees were comprehensive, whereas the PIC/PICA and LIC/LICA scores (IC/ICA scores adjusted under the “Probabilistic” and “Lossless” schemes, respectively) were reported if some evaluation trees were partial. The underlying bipartition frequencies for calculating the IC/ICA scores were obtained by turning on the “-C” option in RAxML. The quartet-based IC scores were calculated using the program *QuartetScores*, and the LQ-IC/QP-IC/EQP-IC scores were always reported regardless of the status of missing taxa in the evaluation tree set.

### Comparing the performance of quartet-based and bipartition-based IC measures

#### Accuracy

We followed Kobert et al. (2016) to define the accuracy of an IC measure as the distance between the IC scores calculated from evaluation tree sets before and after taxon-pruning. The quartet-based and bipartition-based IC scores were first calculated for each of the Original, L1, L2, L3, E1, and E2 data sets. Then, for each type of IC scores and each reference tree, pairwise distances were calculated between the Original data set and each of the five pruned data sets. The accuracy was measured by pairwise distance instead of Spearman’s (or Pearson’s) correlation coefficient because two sets of very different scores can still have very high correlation coefficient (e.g., scores in one set are one tenth of the corresponding scores in the other set). However, unlike in Kobert et al. (2016), here we used the Euclidean distance:

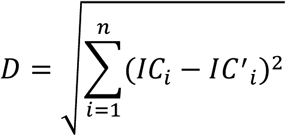

where *n* is the number of internodes in the reference tree (*n* = 98 for these simulated data sets), *IC_i_* and *IC*'_*i*_ refer to the IC scores based on the Original and the pruned data sets, respectively, for the same internode *i*. In addition, in each pairwise comparison, we also calculated the fraction of internodes for which the IC scores were overestimated (by more than 0.05) on the pruned data set compared to the Original data set.

#### Random evaluation tree test

In this test, the topologies of all evaluation trees in the Original, E1, and E2 data sets were randomized. Since the randomized evaluation trees have the same sets of taxa as the original trees, the pattern of missing taxa in each data set was kept the same. The quartet-based and bipartition-based IC scores were then calculated from the original reference trees and randomized evaluation trees.

#### Altered reference tree test

Here, the topologies of the reference trees were altered, whereas the evaluation tree topologies remained unchanged. First, we calculated the relative Robinson-Foulds (rRF) distance (Robinson 1971) between each evaluation tree in the Original data set and its corresponding reference tree. Polytomous evaluation trees were randomly resolved before calculating rRF distances. Second, we classified the evaluation trees in to five categories based on their rRF distances; the ranges of rRF distances of the five categories were: [0.1, 0.3), [0.3, 0.5), [0.5, 0.7), [0.7, 0.9), and [0.9, 1]. Finally, we randomly selected 10 evaluation trees from each category to be the new reference trees. The evaluation tree sets to which the new reference trees belonged were also selected as the new evaluation tree sets. Similarly, for data sets E1 and E2, the evaluation tree sets that match with the new reference trees were selected for this test. The quartet-based and bipartition-based IC scores were then calculated from the new, altered reference trees and their corresponding evaluation trees.

## Acknowledgements

This work was conducted in part using the resources of the Advanced Computing Center for Research and Education at Vanderbilt University. This work was supported by the National Science Foundation (DEB-1442113 to A.R.). Part of this work was financially supported by the Klaus Tschira Foundation.

## References

Aberer AJ, Stamatakis A editors. 2011 IEEE International Conference on Bioinformatics and Biomedicine. 2011 12-15 Nov. 2011.

Anisimova M, Gascuel O. 2006. Approximate likelihood-ratio test for branches: A fast, accurate, and powerful alternative. Systematic Biology 55:539–552.

Anisimova M, Gil M, Dufayard JF, Dessimoz C, Gascuel O. 2011. Survey of branch support methods demonstrates accuracy, power, and robustness of fast likelihood-based approximation schemes. Systematic Biology 60:685–699.

Avni E, Cohen R, Snir S. 2015. Weighted quartets phylogenetics. Systematic Biology 64: 233–242.

Borowiec ML, Lee EK, Chiu JC, Plachetzki DC. 2015. Extracting phylogenetic signal and accounting for bias in whole-genome data sets supports the Ctenophora as sister to remaining Metazoa. BMC Genomics 16:987.

Chen MY, Liang D, Zhang P. 2015. Selecting Question-Specific Genes to Reduce Incongruence in Phylogenomics: A Case Study of Jawed Vertebrate Backbone Phylogeny. Systematic Biology 64: 1104–1120.

Chesters D. 2017. Construction of a Species-Level Tree of Life for the Insects and Utility in Taxonomic Profiling. Systematic Biology 66: 426–439.

Chifman J, Kubatko L. 2014. Quartet inference from SNP data under the coalescent model. Bioinformatics 30: 3317–3324.

Degnan JH, Rosenberg NA. 2009. Gene tree discordance, phylogenetic inference and the multispecies coalescent. Trends in Ecology & Evolution 24: 332–340.

Gadagkar SR, Rosenberg MS, Kumar S. 2005. Inferring species phylogenies from multiple genes: concatenated sequence tree versus consensus gene tree. Journal of Experimental Zoology, Part B: Molecular and Developmental Evolution 304: 64–74.

Guindon S, Dufayard JF, Lefort V, Anisimova M, Hordijk W, Gascuel O. 2010. New algorithms and methods to estimate maximum-likelihood phylogenies: assessing the performance of PhyML 3.0. Systematic Biology 59: 307–321.

Jarvis ED, Mirarab S, Aberer AJ, Li B, Houde P, Li C, Ho SY, Faircloth BC, Nabholz B, Howard JT, et al. 2014. Whole-genome analyses resolve early branches in the tree of life of modern birds. Science 346: 1320–1331.

Jeffroy O, Brinkmann H, Delsuc F, Philippe H. 2006. Phylogenomics: the beginning of incongruence? Trends in Genetics 22: 225–231.

Kobert K, Salichos L, Rokas A, Stamatakis A. 2016. Computing the Internode Certainty and Related Measures from Partial Gene Trees. Molecular Biology and Evolution 33: 1606–1617.

Krabberod AK, Orr RJS, Brate J, Kristensen T, Bjorklund KR, Shalchian-Tabrizi K. 2017. Single Cell Transcriptomics, Mega-Phylogeny, and the Genetic Basis of Morphological Innovations in Rhizaria. Molecular Biology and Evolution 34: 1557–1573.

Kumar S, Filipski AJ, Battistuzzi FU, Kosakovsky Pond SL, Tamura K. 2012. Statistics and truth in phylogenomics. Molecular Biology and Evolution 29: 457–472.

Leveille-Bourret E, Starr JR, Ford BA, Lemmon EM, Lemmon AR. 2017. Resolving Rapid Radiations Within Angiosperm Families Using Anchored Phylogenomics. Systematic Biology.

Li Z, Defoort J, Tasdighian S, Maere S, Van de Peer Y, De Smet R. 2016. Gene Duplicability of Core Genes Is Highly Consistent across All Angiosperms. Plant Cell 28: 326–344.

Maddison WP. 1997. Gene Trees in Species Trees. Systematic Biology 46: 523–536.

Mirarab S, Warnow T. 2015. ASTRAL-II: coalescent-based species tree estimation with many hundreds of taxa and thousands of genes. Bioinformatics 31:i44–52.

Misof B, Liu S, Meusemann K, Peters RS, Donath A, Mayer C, Frandsen PB, Ware J, Flouri T, Beutel RG, et al. 2014. Phylogenomics resolves the timing and pattern of insect evolution. Science 346: 763–767.

Nagy LG, Ohm RA, Kovacs GM, Floudas D, Riley R, Gacser A, Sipiczki M, Davis JM, Doty SL, de Hoog GS, et al. 2014. Latent homology and convergent regulatory evolution underlies the repeated emergence of yeasts. Nature Communications 5:4471.

Nieselt-Struwe K, von Haeseler A. 2001. Quartet-mapping, a generalization of the likelihood-mapping procedure. Molecular Biology and Evolution 18: 1204–1219.

Pease JB, Brown JW, Walker JF, Hinchliff CE, Smith SA. 2017. Quartet Sampling distinguishes lack of support from conflicting support in the plant tree of life. bioRxiv.

Prum RO, Berv JS, Dornburg A, Field DJ, Townsend JP, Lemmon EM, Lemmon AR. 2015. A comprehensive phylogeny of birds (Aves) using targeted next-generation DNA sequencing. Nature 526: 569–573.

Ranwez V, Gascuel O. 2001. Quartet-based phylogenetic inference: improvements and limits. Molecular Biology and Evolution 18: 1103–1116.

Robinson DF. 1971. Comparison of labeled trees with valency three. Journal of Combinatorial Theory, Series B 11: 105–119.

Salichos L, Rokas A. 2013. Inferring ancient divergences requires genes with strong phylogenetic signals. Nature 497: 327–331.

Salichos L, Stamatakis A, Rokas A. 2014. Novel information theory-based measures for quantifying incongruence among phylogenetic trees. Molecular Biology and Evolution 31: 1261–1271.

Sayyari E, Mirarab S. 2016. Fast Coalescent-Based Computation of Local Branch Support from Quartet Frequencies. Molecular Biology and Evolution 33: 1654–1668.

Shannon CE. 1948. A mathematical theory of communication. Bell System Technical Journal 27.

Shen XX, Hittinger CT, Rokas A. 2017. Contentious relationships in phylogenomic studies can be driven by a handful of genes. Nature Ecology & Evolution 1:0126.

Shen XX, Zhou X, Kominek J, Kurtzman CP, Hittinger CT, Rokas A. 2016. Reconstructing the Backbone of the Saccharomycotina Yeast Phylogeny Using Genome-Scale Data. G3: Genes|Genomes|Genetics 6: 3927–3939.

Slowinski JB, Page RD. 1999. How should species phylogenies be inferred from sequence data? Systematic Biology 48: 814–825.

Smith SA, Moore MJ, Brown JW, Yang Y. 2015. Analysis of phylogenomic datasets reveals conflict, concordance, and gene duplications with examples from animals and plants. BMC Evolutionary Biology 15:150.

Song S, Liu L, Edwards SV, Wu S. 2012. Resolving conflict in eutherian mammal phylogeny using phylogenomics and the multispecies coalescent model. Proceedings of the National Academy of Sciences, USA 109: 14942–14947.

Strimmer K, von Haeseler A. 1997. Likelihood-mapping: a simple method to visualize phylogenetic content of a sequence alignment. Proceedings of the National Academy of Sciences, USA 94: 6815–6819.

Strimmer K, von Haeseler A. 1996. Quartet Puzzling: A Quartet Maximum-Likelihood Method for Reconstructing Tree Topologies. Molecular Biology and Evolution 13: 964–964.

Struck TH, Golombek A, Weigert A, Franke FA, Westheide W, Purschke G, Bleidorn C, Halanych KM. 2015. The evolution of annelids reveals two adaptive routes to the interstitial realm. Current Biology 25: 1993–1999.

Tarver JE, Dos Reis M, Mirarab S, Moran RJ, Parker S, O’Reilly JE, King BL, O’Connell MJ, Asher RJ, Warnow T, et al. 2016. The Interrelationships of Placental Mammals and the Limits of Phylogenetic Inference. Genome Biology and Evolution 8: 330–344.

Wang Y, Zhou X, Yang D, Rokas A. 2015. A Genome-Scale Investigation of Incongruence in Culicidae Mosquitoes. Genome Biology and Evolution 7: 3463–3471.

Whelan NV, Kocot KM, Moroz LL, Halanych KM. 2015. Error, signal, and the placement of Ctenophora sister to all other animals. Proceedings of the National Academy of Sciences, USA 112: 5773–5778.

Wickett NJ, Mirarab S, Nguyen N, Warnow T, Carpenter E, Matasci N, Ayyampalayam S, Barker MS, Burleigh JG, Gitzendanner MA, et al. 2014. Phylotranscriptomic analysis of the origin and early diversification of land plants. Proceedings of the National Academy of Sciences, USA 111:E4859–4868.

Wilkinson M. 2006. Identifying stable reference taxa for phylogenetic nomenclature. Zoologica Scripta 35: 109–112.

Xi Z, Liu L, Rest JS, Davis CC. 2014. Coalescent versus concatenation methods and the placement of Amborella as sister to water lilies. Systematic Biology 63: 919–932.

Yang Y, Moore MJ, Brockington SF, Soltis DE, Wong GK, Carpenter EJ, Zhang Y, Chen L, Yan Z, Xie Y, et al. 2015. Dissecting Molecular Evolution in the Highly Diverse Plant Clade Caryophyllales Using Transcriptome Sequencing. Molecular Biology and Evolution 32: 2001–2014.

Zhong B, Liu L, Yan Z, Penny D. 2013. Origin of land plants using the multispecies coalescent model. Trends in Plant Science 18: 492–495.

Zhou X, Shen X-X, Hittinger CT, Rokas A. 2017. Evaluating Fast Maximum Likelihood-Based Phylogenetic Programs Using Empirical Phylogenomic Data Sets. bioRxiv.

